# Reconstruction of genome--scale metabolic model for *Hansenula polymorpha* using RAVEN toolbox

**DOI:** 10.1101/2021.06.18.448943

**Authors:** Francisco Zorrilla, Eduard J Kerkhoven

## Abstract

Genome scale metabolic models (GEMs) provide a useful framework for modeling the metabolism of microorganisms. While the applications of GEMs are wide and far reaching, the reconstruction and continuous curation of such models can be perceived as a tedious and time consuming task. Using the RAVEN toolbox, this protocol practically demonstrates how researchers can create their own GEMs using a homology based approach. To provide a complete example, we reconstruct a draft GEM for the industrially relevant yeast *Hansenula polymorpha*.

## 1 Introduction

Cellular metabolism are complex modular networks involving several hundreds of reactions and metabolites and to meaningfully describe their behaviour requires computational approaches. Genome-scale metabolic models (GEM) are instrumental, as they present a mathematical description of the metabolic network of a given organism in a quantitative manner. Central to the mathematical formulation of GEMs is the *S*-matrix, capturing the stoichiometry of biochemical pathways and metabolic capabilities (**Figure 1**). Computational models such as GEMs are generally based on a bottom-up approach, leveraging the plethora of biochemical and omics information generated by the scientific community to reconstruct the complex network of metabolism. Well curated GEMs can be powerful tools suitable for a large range of analyses. Most notably, constraining the model with measurable reaction rates such as carbon uptake allow for estimation of all metabolic fluxes using flux balance analysis (FBA (Orth et al. 2010)). Therefore, GEMs can be used for the prediction of phenotypic effects caused by genetic modifications. This has a vast range of applications including antibiotic discovery, production strain engineering, and microbial community interaction analysis (Lewis et al. 2012). In short, well curated GEMs are useful frameworks that allow researchers to supplement *in vitro/in vivo* experiments with *in silico* simulations.

**Figure 1.**
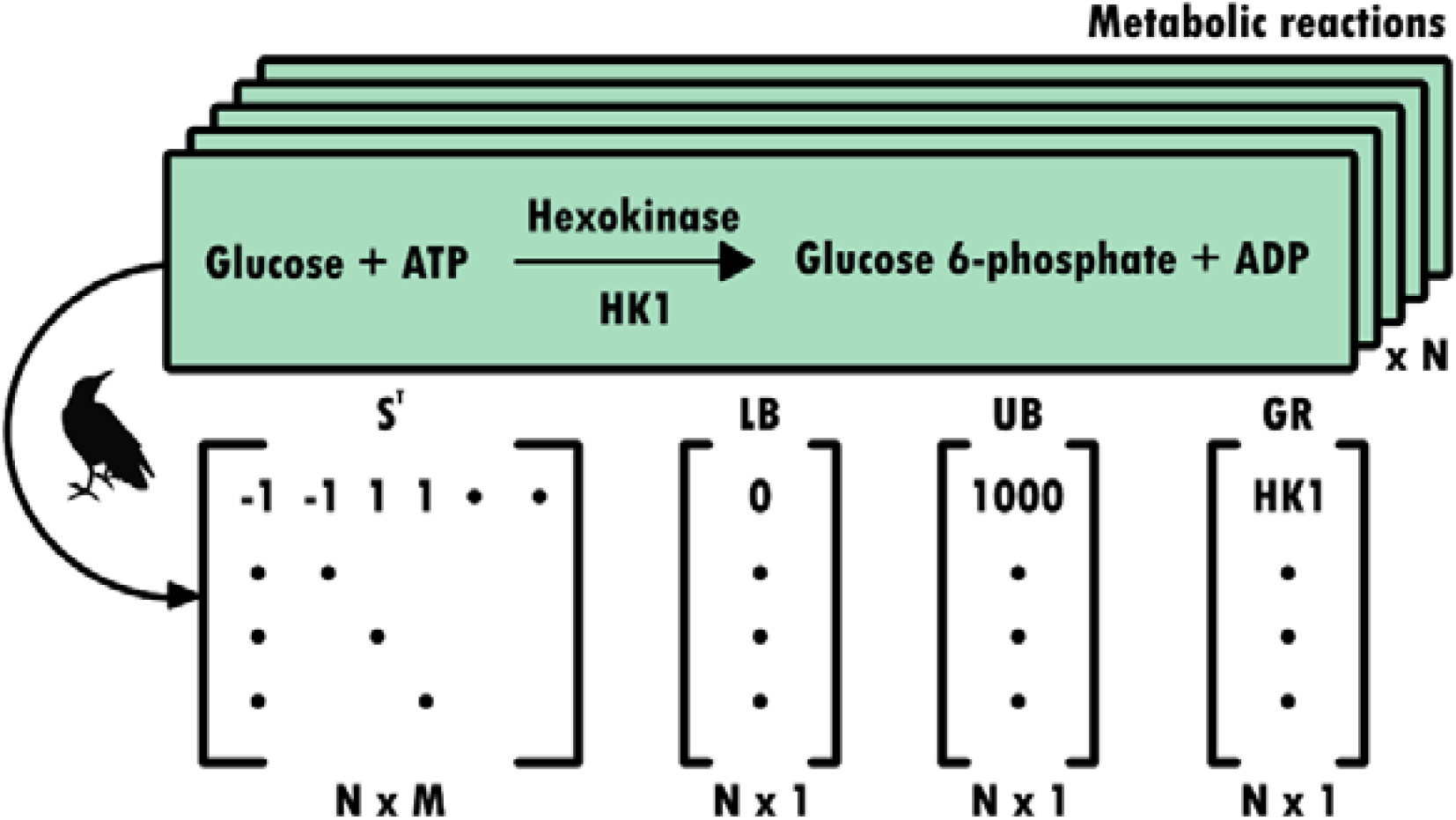
Conceptual representation of the information stored in a genome scale metabolic model. A GEM is fundamentally based on the S-matrix, which is an elegant summary of the stoichiometry of an organism’s specific biochemical pathways and metabolic capabilities. This matrix is sparse and of dimensions M by N, where M represents the number of metabolites and N the number of reactions present in the metabolism of a given organism. Each reaction in this matrix is constrained by some lower bound (LB) and upper bound (UB), which reflects the biological constraints of each reaction (e.g. thermodynamics, reversibility). Additionally, genetic information about each enzyme catalyzed reaction is stored in the grRules structure.

The first published GEM, which debuted in 1999, aimed to model the metabolism of the modest *Haemophilus influenzae Rd*. Since then, GEMs for larger and more complex organisms have been developed and extensively curated. In 2010, the existence of over 30 published GEMs was reported (Thiele and Palsson 2010), while that number has increased significantly since then as the GEM reconstruction process continues to mature. To facilitate the reconstruction, curation and analysis of models, a number of toolboxes have been developed, e.g. COBRA (Heirendt et al. 2017) and RAVEN Toolboxes (Wang et al. 2018). While the reconstruction of GEMs has come a long way since the turn of the millennium, there are a number of challenges that persist in the reconstruction process. For example, manual curation and model refinement represents a glaring bottleneck in the reconstruction process, as this step can take anywhere from weeks to years.

RAVEN can create draft GEMs for an organism of interest by using models from phylogenetically related organisms as templates. This functionality allows users to leverage well curated models to develop GEMs for new organisms of relevance. To reconstruct a GEM for *Hansenula polymorpha*, the highly curated model for the ascomycete *Saccharomyces cerevisiae* will be used as primary template, while a model of the basidiomycete *Rhodosporidium toruloides* is used as complementary template model.

The following protocol demonstrates a practical implementation of the various steps necessary for the homology--based reconstruction of GEMs using the RAVEN toolbox. It assumes a basic knowledge of GEMs and constraint-based modelling, which otherwise are reviewed here (Orth et al. 2010; Thiele and Palsson 2010; Lewis et al. 2012). The approach described in this protocol leverages on high--quality GEMs of phylogenetically related organisms and uses these models as templates to build a new organism-specific model. Whether reactions from the template models should be included is informed through querying homology between the protein sequences of the organisms. As an illustrative example, a draft model for the industrially relevant filamentous yeast *Hansenula polymorpha* (also known as *Ogataea polymorpha*) will be reconstructed from homology with highly curated models. *H. polymorpha* is of scientific interest while lacking a published high-quality GEM. Indeed, this organism was recently demonstrated to be conveniently editable with a CRISPR/Cas9 system (Numamoto et al. 2017), rendering it an even more attractive target for GEM reconstruction.

## 2 Materials

GEM reconstruction requires various software packages (**Table 1**). In addition, several files related to the organism of interest and to the organism(s) that are used as template models are required (**Table 2**).

**Table 1:** Summary of software dependencies and sources.

| Software | Dependencies | Source |
| --- | --- | --- |
| MATLAB | License | https://mathworks.com/products/matlab.html |
| RAVEN | libSBML, Solver | https://github.com/SysBioChalmers/RAVEN |
| COBRA | MATLAB | https://github.com/opencobra/cobratoolbox |
| Gurobi | License | https://www.gurobi.com/products/gurobi-optimizer/ |
| libSBML | MATLAB | http://sbml.org/Software/libSBML/Downloading_libSBML |

**Table 2:** Summary of required organism specific file types.

|  | File Type | Usage | Organism | Source |
| --- | --- | --- | --- | --- |
| Template organism | GEM | Template network | *S. cerevisiae* | https://github.com/SysBioChalmers/yeast-GEM/releases/tag/v8.3.0 |
|  | GEM | Template network | *R. toruloides* | https://github.com/SysBioChalmers/rhto-GEM/releases/tag/v1.1.1 |
|  | Protein FASTA | BLAST | *S. cerevisiae* | https://github.com/SysBioChalmers/rhto-GEM/blob/master/ComplementaryData/genome/sce_s288c.faa |
|  | Protein FASTA | BLAST | *R. toruloides* | https://github.com/SysBioChalmers/rhto-GEM/blob/master/ComplementaryData/genome/rhto_np11.faa |
| Target organism | Protein FASTA | BLAST | *H. polymorpha* | https://genome.jgi.doe.gov/portal/Hanpo2/download/Hanpo2_GeneCatalog_proteins_20100927.aa.fasta.gz |
|  | DNA FASTA | Biomass curation | *H. polymorpha* | ftp://ftp.ncbi.nlm.nih.gov/genomes/all/GCF/001/664/045/GCF_001664045.1_Hanpo2/GCF_001664045.1_Hanpo2_genomic.fna.gz |
|  | Genbank | Biomass curation | *H. polymorpha* | ftp://ftp.ncbi.nlm.nih.gov/genomes/all/GCF/001/664/045/GCF_001664045.1_Hanpo2/GCF_001664045.1_Hanpo2_genomic.gbff.gz |

### 2.1 Software

This protocol is based on the computing environment MATLAB, complemented with software specific for reconstructing and analysing genome-scale models. Note that most software is open source and available through GitHub, or offers an academic license.

1. Although not open source, MATLAB is a widely used programming language in science and engineering. A MATLAB license can be purchased privately or obtained through an academic institution. This protocol requires version 2013b or higher, no additional MathWorks toolboxes are required.
2. RAVEN, for Reconstruction, Analysis and Visualization of Metabolic Networks, is an open source MATLAB toolbox for GEM development and analysis. RAVEN is freely available (Table 1), works on Windows, Mac and Unix, and has an active user and developer community (*see* **Note 1**).
3. To facilitate import and export of GEMs, RAVEN depends on the libSBML MATLAB API that facilitates handling of the community standard SBML file format. libSBML is freely available (**Table 1**) and installation descriptions are provided.
4. Simulation of GEMs depends on a linear programming solver. There are various solvers that can interface with MATLAB, the protocol uses Gurobi, which is available under an academic license (**Table 1**). Alternatively, RAVEN can utilize the open source glpk solver that is part of a COBRA installation (Heirendt et al. 2017).

### 2.2 Files

To reconstruct a new GEM requires various files related to the organism of interest. In addition, homology--based reconstruction requires files related to the high-quality models that will be used as templates (**Table 2, Figure 3**).

**Figure 2.**
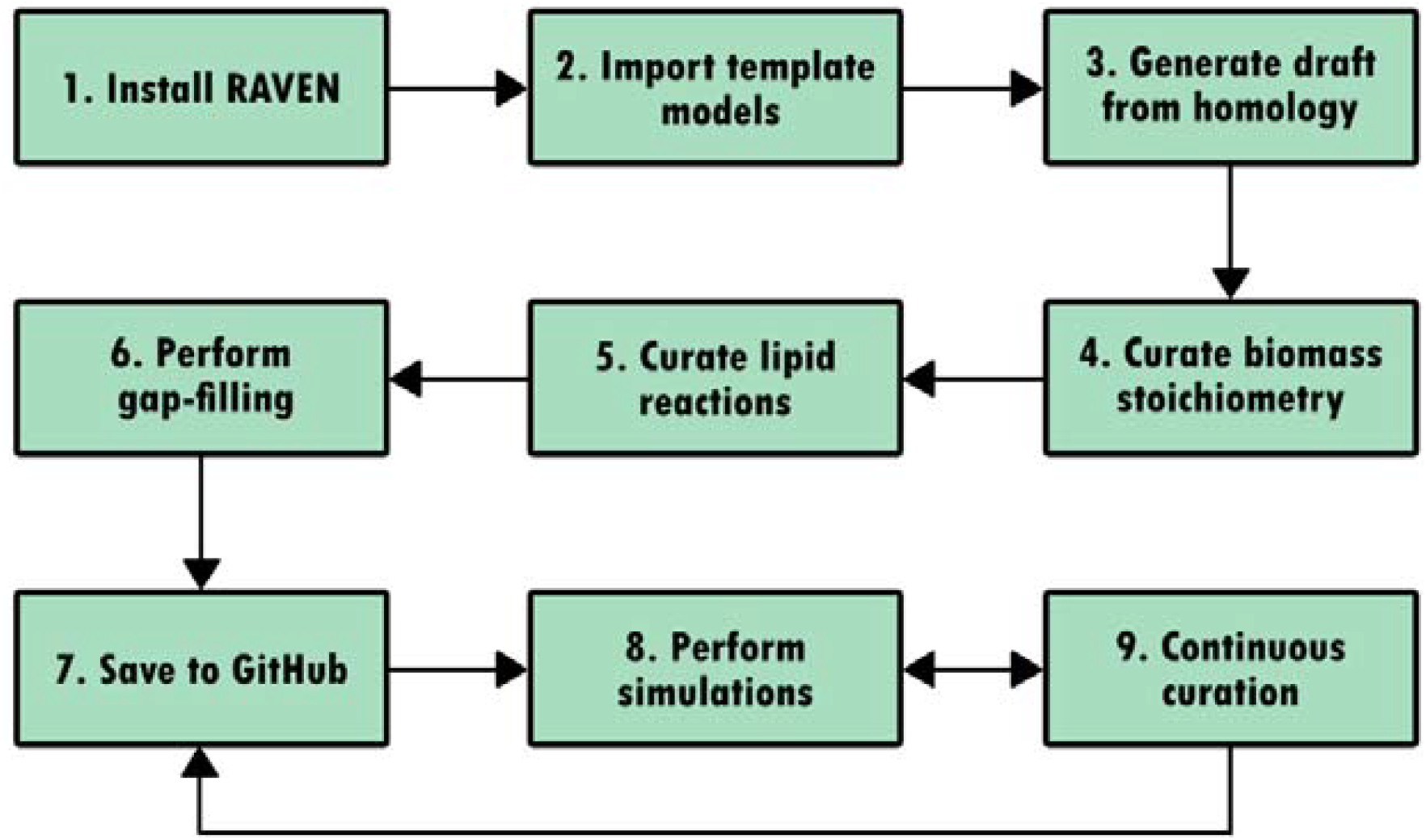
Flow diagram representation of this protocol. Note that each box corresponds to a subsection in the Methods section.

**Figure 3.**
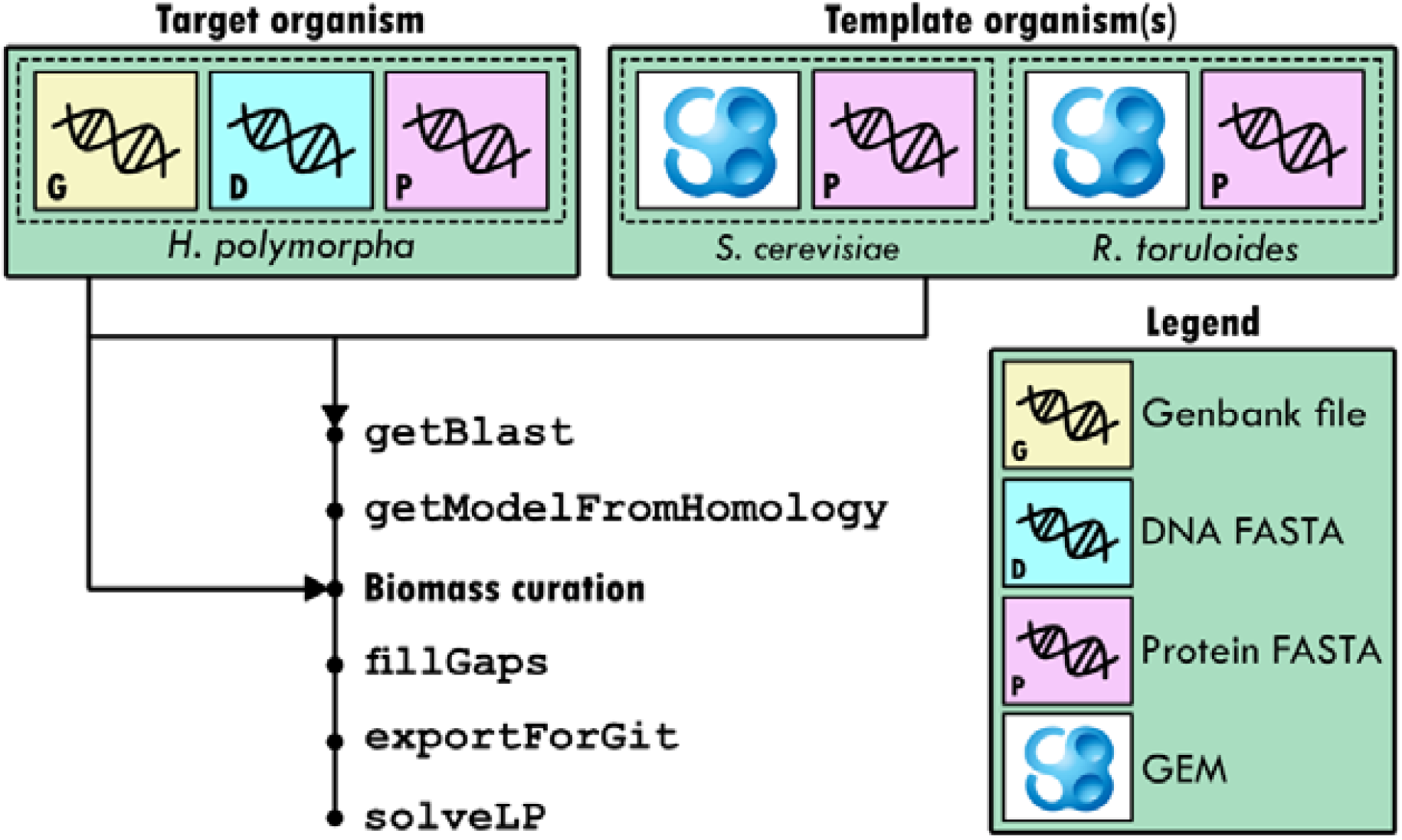
Visual summary of all the files discussed in Table 2. Various key RAVEN functions are shown in Courier New font. Arrows show which files are used by the different functions and tasks. Each dot in the flow diagram represents an intermediate version of the target GEM.

1. For the organism of interest: (1) protein FASTA with sequences of all proteins in its genome; (2) DNA FASTA for determining nucleotide ratios, and (3) Genbank file to determine ribonucleotide ratios. These files can be obtained individually from their respective original sources. For this protocol, files for *H. polymorpha* are provided on the GitHub repository.
2. For template organism(s): (1) protein FASTA with sequences of all proteins in its genome; (2) model file in SBML format. Important for automatic matching of is that the protein FASTA and the model file use the same gene identifier (*see* **Note 2**).

### 2.3 Literature data

The model can be curated, and its quality improved, leveraging on *any* available knowledge on the metabolic behaviour and capabilities of the organism of interest. This includes the levels of macromolecules that make up biomass; knowledge on essential and non-essential genes; the ability to utilize different nutrient sources; and growth rate during cultivation. Through a brief literature search, literature reporting total protein content, lipid and carbohydrate levels can be identified for *H. polymorpha* (Petersen 1985; Wijeyaratne et al. 1986), which can be used to specify the biomass reactions (section 3.4). Data from related organism can be used to estimate macromolecule levels in lieu of organism specific data.

## 3 Methods

The reconstruction of the *H. polymorpha* GEM is tracked in a git repository as detailed in section 3.7. All required files and scripts are hosted online at GitHub (*see* **Note 3**), where hanpo-GEM is used as a convenient short name for the model. Download (or *clone*, in git nomenclature) the hanpo-GEM repository to obtain a local copy of all the content. The `reconstructionProtocol` script contains all required commands to obtain a functional draft model of *H. polymorpha* metabolism: hanpo-GEM. Note that the commands given in this chapter, recognizable by `font`, are occasionally more concise than in the script, typically related to the location of files in the repository. An overview of the various steps in GEM reconstruction is shown in **Figure 2** and discussed in detail below.

### 3.1 Install RAVEN

After obtaining all required software and files (see Methods), the relevant software needs to be installed. Detailed installation instructions, requirements and dependencies are detailed on the RAVEN Wiki (*see* **Note 1**). To verify that RAVEN and its dependencies been properly installed, run the `checkInstallation` function in the MATLAB command window. The output of this function is useful for troubleshooting installation problems.

### 3.2 Import template models

RAVEN can create draft GEMs for an organism of interest by using models from phylogenetically related organisms as templates (an alternative approach is *de novo* reconstruction, *see* **Note 4**). This functionality allows users to leverage well curated models to develop GEMs for new organisms of relevance. To reconstruct a GEM for *H. polymorpha*, the highly curated model for the ascomycete *Saccharomyces cerevisiae* will be used as primary template, while a model of the basidiomycete *Rhodosporidium toruloides* is used as complementary template model. While there is no limit to the number of template models one can use, it is essential that the model components (i.e. metabolites, reactions, genes) use the same identifiers.

To initiate the reconstruction process, we import the *S. cerevisiae* model into the MATLAB environment with the command:

~~~
modelSce = importModel(‘yeastGEM.xml’);
~~~

The `importModel` function loads the SBML file (with .xml extension) containing the model into a RAVEN format MATLAB structure. The model can now be queried from within MATLAB as `modelSce`, and as novel user it can be informative to inspect its content. For a more user-friendly format to inspect the model, it can be exported as a spreadsheet:

~~~
exportToExcelFormat(modelSce, ‘modelSce.xlsx’);
~~~

This function takes two input arguments: the model structure and a character string of the desired name for the spreadsheet file. The generated Excel file has five sheets: RXNS, METS, COMPS, GENES, and MODEL, contain information regarding each reaction, metabolite, compartment, gene, and model, respectively. While convenient for inspection, users are discouraged to modify the Excel file, but rather make any desired changes to the model in MATLAB and export in SBML format when desired:

~~~
exportModel(modelSce, ‘modelSce.xml’);
~~~

In the reconstruction script similar commands are used to load the *R. toruloides* model.

### 3.3 Generate models from homology

The protein FASTA files from the target and template organisms will be queried to facilitate constructing a draft model for *H. polymorpha* based on homology. The RAVEN functions `getBlast` and `getModelFromHomology` are respectively used for alignment of the protein sequences and subsequent use of these results to reconstruct the first model of the target organism.

#### 3.3.1 Clean and match protein identifiers

It is essential that the gene identifiers in the template model and the corresponding FASTA file match. This can be checked by opening the FASTA files using a text editor and comparing the protein identifier format found here with the gene identifiers in the respective template models (c.f. *S. cerevisiae* YML001W). For the target organism it is also convenient if the protein identifiers are short and corresponding to a format commonly used for that organism. The *H. polymorpha* protein FASTA downloaded here (**Table 2**) requires some editing of the identifiers to remove unnecessary annotations, to yield a convenient format such as *Hanpo2_12345*. This can be accomplished through many different approaches (*see* **Note 2**).

#### 3.3.2 First draft model based on homology

Evidence of homology between target and template protein sequences are determined by bi-directional BLAST:

~~~
blast = getBlast(‘hanpo’, ‘hanpo.faa’, {‘sce’, ‘rhto’}, {‘sce.faa’,
                                        ‘rhto.faa’});
~~~

The `getBlast` function takes four input arguments: a string with the target organism model identifier (‘hanpo’), a string specifying the protein FASTA filename of the target organism (‘hanpo.faa’), a cell containing the strings of the template organism identifiers ({‘sce’,’rhto’}), and a cell containing the strings specifying the protein FASTA filenames of the template organisms ({‘sce.faa’,’rhto_np11.faa’}). This process can take several minutes, and the results are stored in the `blast` variable.

Subsequently, the `blast` variable is used to generate the first draft model for *H. polymorpha* based on homology with *S. cerevisiae* and *R. toruloides*:

~~~
model = getModelFromHomology({modelSce, modelRhto}, blast, ‘hanpo’,
                      {‘sce’, ‘rhto’});
~~~

Multiple cut-offs can be set to affect whether a BLAST hit between two proteins is indicative of homology and supports presence of the corresponding reaction in the target model (*see* **Note 6**).

The homology-based approach is only able to add reactions to the target draft model that are gene annotated in the template models. Exchange reactions are not annotated to genes, as they do not represent enzymatic reactions but rather simulate the flow of metabolites in and out of the modelled system, representing e.g. the depletion of carbon source and accumulation of excreted nutrient. To allow the target model to consume the required nutrients and excrete e.g. CO_2_ we add the required exchange reactions corresponding to growth medium components:

~~~
mediumComps = {‘r_1654’, ‘r_1672’, ‘r_1808’, ‘r_1832’, ‘r_1861’,
          ‘r_1992’, ‘r_2005’, ‘r_2060’, ‘r_2100’, ‘r_2111’};
      model = addRxnsGenesMets(model,modelSce,mediumComps);
~~~

Here the exchange reactions represent ammonium; CO_2_; glycerol, H^+^; iron; O_2_; phosphate; sulphate; water and biomass that should be able to either exchange in or out of the system to support growth.

### 3.4 Define biomass composition

One of the essentialities of metabolism is that it can synthesize the macromolecules that make up the biomass and thereby support the growth of the cell (or, cell division). GEMs contain reactions specifying which macromolecules (e.g. DNA, lipids) are required in which amount, and to obtain an organism-specific GEM requires modifying these reactions to represent the target organism. Defining this at a relatively early stage of the reconstruction ensures that the model can biosynthesize all required macromolecules. Defining the biomass composition requires experimental data such as the dry cell weight fractions of macromolecular components, the CTGA content of its genome, and profile of lipid classes. One typically first determines the levels of each grouping of biomass constituents, ie. DNA, RNA, protein, lipid and carbohydrate. When these values are set, by measurement or inferred from published data, the level of their constitutive parts should be determined.

In the *S. cerevisiae* and *R. toruloides* GEMs, the biomass composition is split over several pseudoreactions, such as the DNA pseudoreaction:

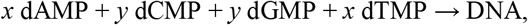

while all macromolecules are then combined to generate biomass:

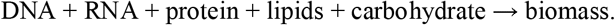

For the target model, start by taking the biomass pseudoreactions from a template model, which can here be easily recognized by a reaction name ending with *pseudoreaction*:

~~~
biomassRxns = endsWith(modelRhto.rxnNames, ‘pseudoreaction’);
            biomassRxns = modelRhto.rxns(biomassRxns);
     model = addRxnsGenesMets(model, modelRhto, biomassRxns);
~~~

A list of their reaction identifiers is generated, and this is subsequently used to add the relevant reactions from `modelRhto` to `model`. A spreadsheet is provided in the `ComplementaryData/biomass` folder of the model repository detailing calculations required to obtain the stochiometric coefficients of the biomass components for the target organism, while the principles of these calculations are detailed below.

The stoichiometric coefficients, or concentrations of macromolecule constituents, are collated and subsequently used to update the biomass composition. The stoichiometric coefficients of the DNA pseudoreaction can directly be modified as:

~~~
DNA.mets = {‘s_0584’, ‘s_0589’, ‘s_0615’, ‘s_0649’, ‘s_3720’};
DNA.stoichCoeffs = [-0.00189, -0.00174, -0.00174, -0.00189, 1];
     model = changeRxns(model, ‘r_4050’, DNA, 1);
~~~

#### 3.4.1 Nucleotides

To estimate the required level of deoxyribonucleotides that constitute the DNA in the cell, we can use a DNA FASTA file of the *H. polymorpha* genome to determine the frequency of each nucleotide in the genome of our target organism. This can readily be done using `sed` in Unix or similar commands on other systems:

~~~
      sed -i ‘s/>.*$//’ hanpo.fna
 less hanpo.fa | grep a -oi|wc -l
~~~

Where the first line removes the gene identifiers from the FASTA file, and the second line counts the occurrence of each nucleotide. In the template yeast-GEM, a DNA content of 0.3 % of biomass was assumed, which translates to 0.24% for *H. polymorpha* due to its smaller genome size. In addition to considering the total number of nucleotides, the paired nature of CG and AT nucleotides and the molecular weight of each nucleotide allows for estimation of how much deoxyribonucleotides are required to produce 1 g of biomass (**Table 3**).

**Table 3.**
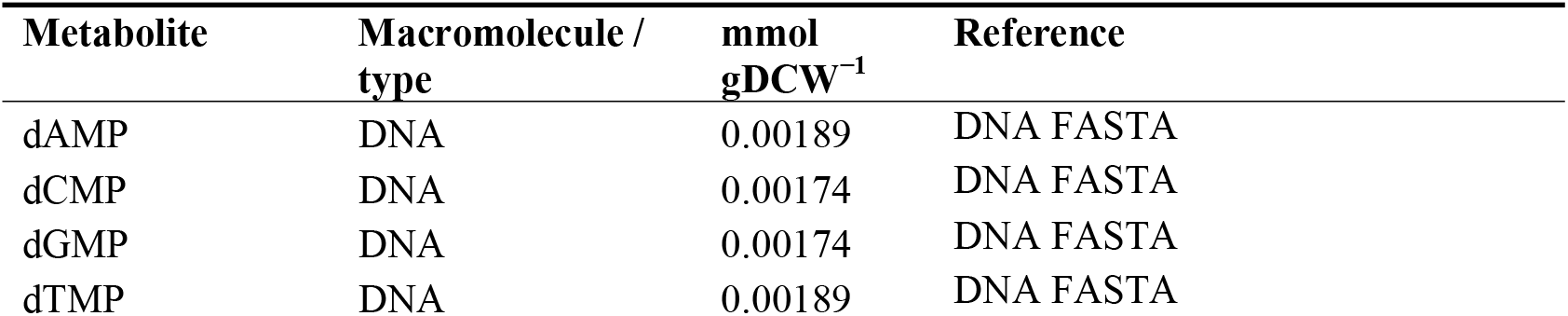
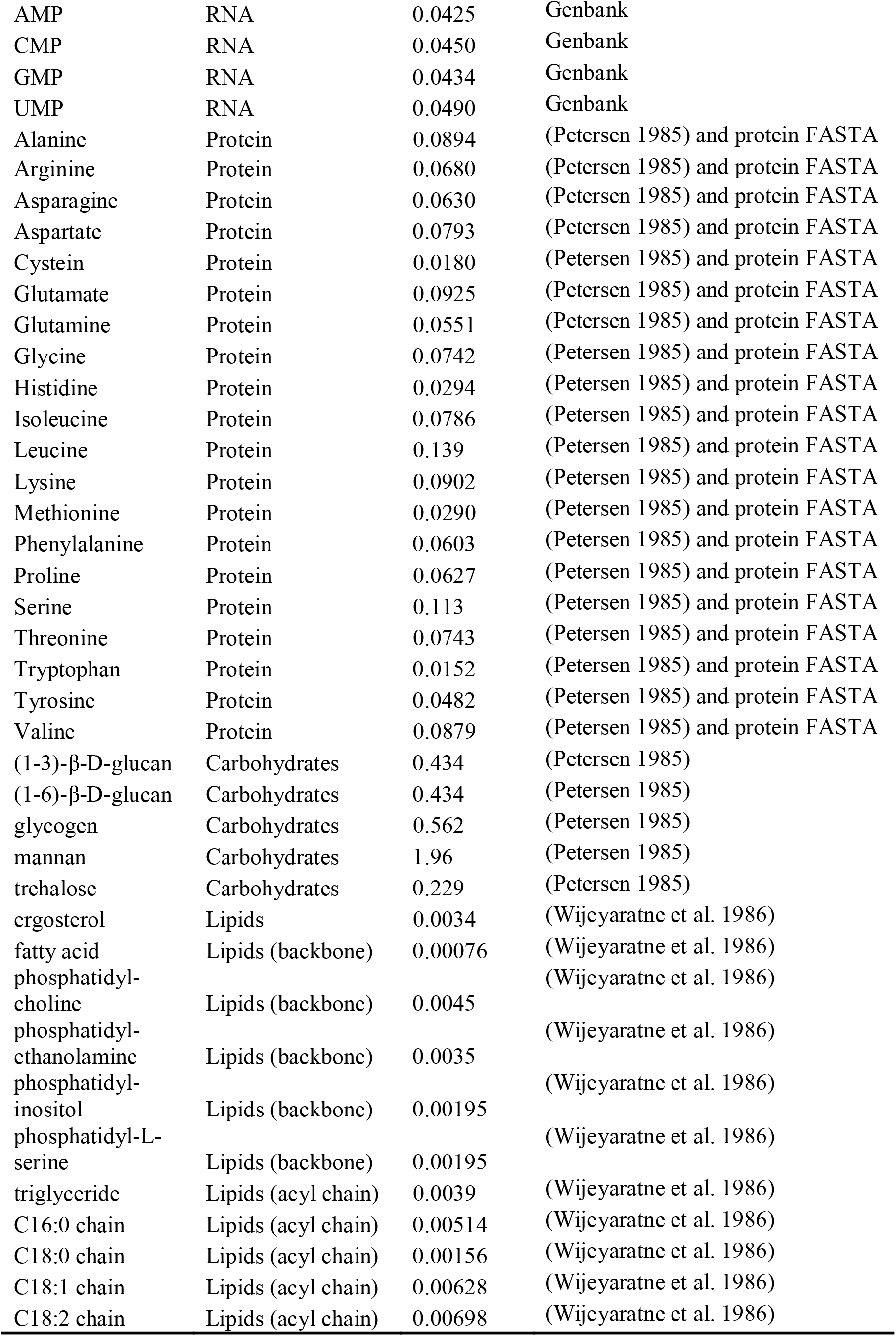
Biomass components.

| Metabolite | Macromolecule / type | mmol gDCW<sup>-1</sup> | Reference |
| --- | --- | --- | --- |
| dAMP | DNA | 0.00189 | DNA FASTA |
| dCMP | DNA | 0.00174 | DNA FASTA |
| dGMP | DNA | 0.00174 | DNA FASTA |
| dTMP | DNA | 0.00189 | DNA FASTA |
| AMP | RNA | 0.0425 | Genbank |
| CMP | RNA | 0.0450 | Genbank |
| GMP | RNA | 0.0434 | Genbank |
| UMP | RNA | 0.0490 | Genbank |
| Alanine | Protein | 0.0894 | (Petersen 1985) and protein FASTA |
| Arginine | Protein | 0.0680 | (Petersen 1985) and protein FASTA |
| Asparagine | Protein | 0.0630 | (Petersen 1985) and protein FASTA |
| Aspartate | Protein | 0.0793 | (Petersen 1985) and protein FASTA |
| Cysteine | Protein | 0.0180 | (Petersen 1985) and protein FASTA |
| Glutamate | Protein | 0.0925 | (Petersen 1985) and protein FASTA |
| Glutamine | Protein | 0.0551 | (Petersen 1985) and protein FASTA |
| Glycine | Protein | 0.0742 | (Petersen 1985) and protein FASTA |
| Histidine | Protein | 0.0294 | (Petersen 1985) and protein FASTA |
| Isoleucine | Protein | 0.0786 | (Petersen 1985) and protein FASTA |
| Leucine | Protein | 0.139 | (Petersen 1985) and protein FASTA |
| Lysine | Protein | 0.0902 | (Petersen 1985) and protein FASTA |
| Methionine | Protein | 0.0290 | (Petersen 1985) and protein FASTA |
| Phenylalanine | Protein | 0.0603 | (Petersen 1985) and protein FASTA |
| Proline | Protein | 0.0627 | (Petersen 1985) and protein FASTA |
| Serine | Protein | 0.113 | (Petersen 1985) and protein FASTA |
| Threonine | Protein | 0.0743 | (Petersen 1985) and protein FASTA |
| Tryptophan | Protein | 0.0152 | (Petersen 1985) and protein FASTA |
| Tyrosine | Protein | 0.0482 | (Petersen 1985) and protein FASTA |
| Valine | Protein | 0.0879 | (Petersen 1985) and protein FASTA |
| (1-3)-β-D-glucan | Carbohydrates | 0.434 | (Petersen 1985) |
| (1-6)-β-D-glucan | Carbohydrates | 0.434 | (Petersen 1985) |
| glycogen | Carbohydrates | 0.562 | (Petersen 1985) |
| mannan | Carbohydrates | 1.96 | (Petersen 1985) |
| trehalose | Carbohydrates | 0.229 | (Petersen 1985) |
| ergosterol | Lipids | 0.0034 | (Wijeyaratne et al. 1986) |
| fatty acid | Lipids (backbone) | 0.00076 | (Wijeyaratne et al. 1986) |
| phosphatidyl-<br>choline | Lipids (backbone) | 0.0045 | (Wijeyaratne et al. 1986) |
| phosphatidyl-<br>ethanolamine | Lipids (backbone) | 0.0035 | (Wijeyaratne et al. 1986) |
| phosphatidyl-<br>inositol | Lipids (backbone) | 0.00195 | (Wijeyaratne et al. 1986) |
| phosphatidyl-L-serine | Lipids (backbone) | 0.00195 | (Wijeyaratne et al. 1986) |
| triglyceride | Lipids (acyl chain) | 0.0039 | (Wijeyaratne et al. 1986) |
| C16:0 chain | Lipids (acyl chain) | 0.00514 | (Wijeyaratne et al. 1986) |
| C18:0 chain | Lipids (acyl chain) | 0.00156 | (Wijeyaratne et al. 1986) |
| C18:1 chain | Lipids (acyl chain) | 0.00628 | (Wijeyaratne et al. 1986) |
| C18:2 chain | Lipids (acyl chain) | 0.00698 | (Wijeyaratne et al. 1986) |

A similar approach can estimate the level of ribonucleotides that constitute RNA. For simplicity, assume that total RNA corresponds to 6% of biomass as was defined in yeast-GEM, and ignore the effects of differential expression of individual RNAs (*see* **Note 7**). Instead, estimate the ribonucleotide distribution on the coding sequences that are annotated in the *H. polymorpha* genome. This information can be obtained from a FASTA file containing only the coding sequences from the genome. Such a file can be generated from a Genbank file and a FASTA parser such as https://rocaplab.ocean.washington.edu/tools/genbank_to_fasta/. The ribonucleotide stoichiometric coefficients are subsequently calculated as described above for deoxyribonucleotides (**Table 3**).

#### 3.4.2 Protein

Also the determination of the stoichiometric coefficients of amino acids for the biomass reactions is a combination of measured total protein content, and determination of the ratio of the constitutive amino acids. Analogous to the nucleotide determination above, amino acid ratios can be determined from the protein FASTA (*see* **Note 7**). These ratios, together with a experimentally measured total protein concentration—obtained from (Petersen 1985) for *H. polymorpha*—are used to determine the stoichiometric coefficients of the amino acids in the biomass reactions (**Table 3**).

#### 3.4.3 Carbohydrates

If experimentally measured, the levels of storage carbohydrates, e.g. glycogen or trehalose, can readily be integrated. While many carbon storage molecules are complex polymers, the reactions in GEMs often only describe the polymerization of one subunit, i.e.

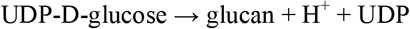

while a more accurate description would be

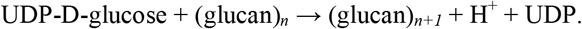

However, GEMs are not able to handle such *groups* of polymers with various lengths. The effect of simplifying the glucan polymerization as above is that the glucan in the model represents one subunit, with chemical formula C_6_H_10_O_5_. This is of relevance when converting glucan levels that are reported in grams to mmol. For *H. polymorpha*, glucan, trehalose and mannan levels are reported (Petersen 1985) and can therefore be incorporated in hanpo-GEM (**Table 3**).

#### 3.4.4 Lipids

Representation of lipid levels in the biomass reaction is typically dependent on two types of quantitative measurements: lipid classes (e.g. triacylglycerol, free fatty acid, phosphatidylinositol) and acyl chains (e.g. 16:0, 16:1, 18:0). There are various ways of representing lipid metabolism, which are discussed in detail here (Kerkhoven 2019). For the *H. polymorpha* model, the SLIME approach is used, which Splits Lipids Into Measurable Entities (Sánchez et al. 2019), as is briefly detailed below. Experimental measurements of lipid classes and acyl chains are reported for *H. polymorpha* (Wijeyaratne et al. 1986), such that the stoichiometric coefficients can be defined (**Table 3**).

### 3.5 Curation of lipid reactions

After adjusting the biomass composition in the model, it might be relevant to curate some reactions that are already known to be different in your organism of interest. This step can also be done *after* gap-filling (detailed below), but it is possible that the curation influences the results from the gap-filling algorithm. For the *H. polymorpha* model, the use of the SLIME formalism to describe lipid metabolism requires curation of the reactions to match the lipids specified in the biomass reactions. The SLIME formalism represents the flexibility of lipid metabolism while allowing incorporation of measurements of lipid classes and acyl chains. While explained in more detail in (Sánchez et al. 2019), briefly, each lipid species that is part of the biomass composition is split into pseudometabolites representing the backbone and acyl chains, which subsequently are gathered to represent the measured entities:

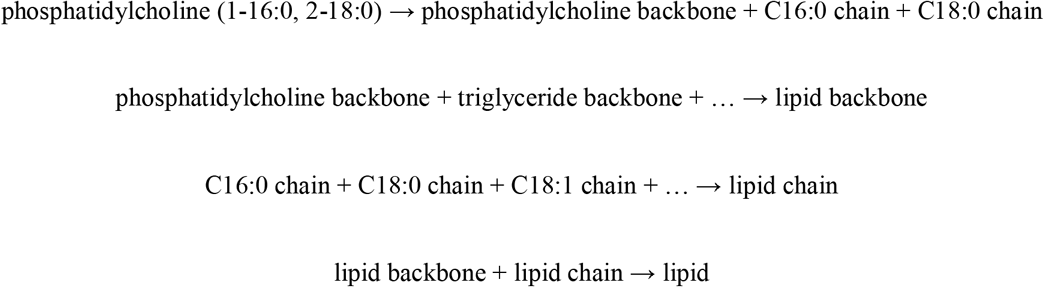

To semi-automatically curate these reactions to match the measured acyl chains and lipid classes, a set of template reactions are provided in the `ComplementaryData/reconstruction` folder of the repository, which are applied to the target model with:

~~~
model = addLipidReactions(template, model);
model = addSLIMEreactions(template, model);
~~~

The target model has now the lipid reactions that are required to support a flux towards lipid biosynthesis. After gap-filling (next section), it becomes apparent that the provided acyl chain and lipid class levels are not fully compatible, and one of the two have to be scaled to yield a functional model. The acyl chain data can be scaled when assuming that the lipid class measurements are technically more reliable:

~~~
model = scaleLipids(model, ‘tails’);
~~~

### 3.6 Perform gap-filling

Homology based draft models tend to have numerous gaps within their metabolic network. These gaps can be thought of as missing reactions in a metabolic pathway required for producing biomass macromolecules. While the pathway is likely functional in the template model, an automated approach such as `getBlast` and `getModelFromHomology` cannot always reliably identify homologs for every gene in this pathway, resulting in incomplete pathways in the draft model. To convert a draft model into a functional model requires identification and resolving of this through gap-filling.

Gaps can be identified and curated through a semi-automatic approach (*see* **Note 8**), however, RAVEN also provides automated gap-filling functionality through the `fillGaps` function, which adds reactions from the template models as required. Here, gaps are resolved to support the use of glycerol as carbon source and biomass production as objective. We therefore force the model to have at non-zero flux through the biomass equation and allow uptake of glycerol as carbon source:

~~~
model = setParam(model, ‘lb’, ‘r_4041’, 0.01);
model = setParam(model, ‘lb’, ‘r_1808’, -10);
~~~

Gaps should be resolved by adding enzymatic and transport reactions from the template models, and not by adding exchange reactions (this could e.g. introduce the use of other carbon sources, which is not satisfactory when gap-filling for growth on glycerol):

~~~
exchangeRxns = getExchangeRxns(modelSce,’both’);
modelSce = removeReactions(modelSce, exchangeRxns);
~~~

The `fillGaps` is provided with the necessary models and subsequently calculates the smallest set of reactions that should be added from the template models to support a flux through the draft model:

~~~
[∼,∼, addedRxns, model, ∼] = fillGaps(model, {modelSce, modelRhto},
 false, true);
~~~

where setting `allowNetProduction` to `false` prevents additional exchange reactions and setting `useModelConstraints` as `true` ensures that the flux from glycerol to biomass formation is gap-filled. This has now generated a model that is able to produce all the macromolecules that constitute the biomass of *H. polymorpha*.

### 3.7 Save to GitHub

To keep track of development and facilitate distribution of the model it is stored in an online GitHub repository (*see* **Note 3**). Every time a significant curation is performed on the model, it should ideally be tracked and committed to the repository. To simplify and standardize this process, functions are provided:

~~~
newCommit(model);
~~~

automatically generates the XML, YML and text files that can aid in identifying differences between model version. This function should be run when a model curation is committed to the repository.

After numerous curations, one might want to designate the existing version of the model as a new numbered release with:

~~~
newRelease(model);
~~~

This function also generates MAT and XLSX model files and ensure that changes between different versions of the model are detailed in the history file. Only when new commits and releases are pushed towards the online GitHub repository do they become publicly available.

### 3.8 Perform simulations

The generated draft model is now able to simulate the production of biomass, which is essential for any living organism, as demonstrated by setting biomass formation as objective function, run flux balance analysis and inspect the active fluxes:

~~~
 model = setParam(model, obj, ‘r_4041’, 1);
 sol = solveLP(model, 1);
 printFluxes(model, sol.x, false)
~~~

When available, the model can be constrained by any measured uptake or excretion of metabolites, by setting the upper or lower bound of the relevant exchange reaction. Through such a process, experimental data reported in literature can be used to evaluate the model and to direct where further development of the model is required.

### 3.9 Manual curation

Once a draft GEM is generated, manual curation is required to obtain a high-quality model that can extensively be used for various types of investigations. The GEM of the model yeast *S. cerevisiae* has been subjected to continuous curation since its first publication in 2003. Manual curation can involve correcting gene associations, removal or addition of reactions, moving reactions to different compartments, while all these curations aim to reach a more accurate description of the *in vivo* metabolic network.

The gap-filling (section 3.6) added reactions from the template models without searching for homology of the associated genes. The draft model therefore currently contains genes from both *S. cerevisiae* and *R. toruloides* that should replaced by their *H. polymorpha* homologs. We can list the reactions that contain a *R. toruloides* gene:

~~~
rhtoRxns = contains(model.grRules,’RHTO’);
~~~

Through manual search in literature and use of BLAST we identify that RHTO_03911, coding for oleoyl-CoA desaturase (reaction y300009), has a homolog in *H. polymorpha*. The gene association of this reaction can therefore be curated:

~~~
model = changeGrRules(model, ‘y300009’, ‘Hanpo2_15704’, true);
~~~

Another curation that can be performed on this model is that *H. polymorpha* is a known methanogen, able to assimilate methanol (Egli et al. 1986). However, the model currently lacks the required reactions. A combination of copying existing reactions from the template model and describing new reactions is sufficient to implement methanol assimilation in the draft model:

~~~
      rxnsToAdd.rxns = {‘MOX’, ‘DAS’};
rxnsToAdd.equations = {‘methanol[p] + oxygen[p] => formaldehyde[p] +
hydrogen peroxide[p]’, ‘formaldehyde[p] + D-xylulose 5-phosphate[p]
         => glyceraldehyde 3-phosphate[p] + glycerone[p]’};
   rxnsToAdd.rxnNames = {‘methanol oxidase’,’dihydroxyacetone
                                 synthase’};
                           rxnsToAdd.lb = [0,0];
                         rxnsToAdd.ub = [1000,1000];
                   rxnsToAdd.eccodes = {‘1.1.3.13’,’2.2.1.3’};
rxnsToAdd.grRules = {‘Hanpo2_76277’,’Hanpo2_95557’};
rxnsToAdd.rxnNotes = {‘Methanol metabolism reaction added by manual curation’,
‘Methanol metabolism reaction added by manual curation’};
model = addRxns(model,rxnsToAdd,3,’’,true,true);
~~~

Where first a `rxnsToAdd` structure is constructed with all relevant information on new reactions, that are subsequently added to the model with the `addRxns` function. In addition with a number of transport and exchange reactions renders the draft model methanogenic, in support of literature.

## 4 Notes

### 4.1 RAVEN documentation and support

With the extensive functionality of RAVEN Toolbox, users might find it useful to consult the documentation that can be found in the `docs` folder. In addition, the RAVEN repository on GitHub has a Wiki with useful information, while the Issues section can be used to report bugs. As example, the Text Analytics Toolbox of MATLAB 2017b and more recent versions has been identified as conflicting with RAVEN Toolbox and a workaround is provided here https://github.com/SysBioChalmers/RAVEN/issues/55. Another channel for support is the Gitter chat forum available at https://gitter.im/SysBioChalmers/RAVEN.

### 4.2 Modifying protein identifiers using sed

It is essential to ensure that protein identifiers match between the model and FASTA file of each organism. One of many ways of accomplishing this is to edit the protein FASTA files using the `sed` command in Unix. The original *S. cerevisiae* protein FASTA contained unwanted annotations after each protein identifiers, that could be readily removed by discarding everything after the first whitespace in every protein identifier:

~~~
sed -i ‘s/ .*$//’ sce.faa
~~~

This replaces instances such as *>YAL001C TFC3 SGDID:S000000001, Chr I from 151006-147594,151166-151097, Genome Release 64-1-1, reverse complement, Verified ORF, “Largest of six subunits of the RNA polymerase III transcription initiation factor complex (TFIIIC); part of the TauB domain of TFIIIC that binds DNA at the BoxB promoter sites of tRNA and similar genes; cooperates with Tfc6p in DNA binding”*, with the corresponding protein identifier, in this case simply:

*>YAL001C*.

### 4.3 hanpo-GEM GitHub repository

A dedicated GitHub repository is used to track the development of hanpo--GEM, available from https://github.com/SysBioChalmers/hanpo-GEM. The repository follows a standardized structure, where `ModelFiles` contains the latest version of hanpo--GEM in various file formats. The `ComplementaryScripts` and `ComplementaryData` folders contain what is required to regenerate the model and run analyses.

### 4.4 *De novo* model generation and model merge

Besides making model from homology, RAVEN also supports *de novo* reconstruction using KEGG or MetaCyc as reaction databases. The functions `getKEGGModelForOrganism` and `getMetacycModelForOrganism` only require the protein FASTA of the organism of interest. This method is beneficial to find reactions that are not part of any template model and can therefore be used as a complementary approach. While *de novo* models can be easily generated, merging them with homology-based models is challenging. Matching of metabolites across the different models is crucial, which is typically based on metabolite identifier or name. However, each database and template model can use their own namespace (*see* **Note 5**). Matching by annotations, such as ChEBI, is a promising alternative approach, however, such databases typically do not cover all metabolites and the models are not always fully annotated. The MNXref namespace attempts to collate and cross-reference between a range of chemical and metabolite databases, but this also suffers from redundancy and duplication (Moretti et al. 2016). To date, no fully automated approaches can therefore be leveraged on, and merging models from different sources will remain to require manual inspection and curation.

### 4.5 Identifiers for model content

Every reaction, metabolite and compartment has a unique identifier in the model (c.f. `model.rxns`). When using multiple models, it is important that the identifiers are defined in the same format. If e.g. glucose 6-phosphate is known in one model as `glc6p`, while `G6P` in another, and `s_0132` in a third model, they will be treated as independent metabolites. When using multiple template models, as demonstrated here, requires that all templates are using the same identifier *namespace*. They do not necessarily have to be informative (cf. `glc6p` and `s_0132`), but they have to be unique and universal for all models used. The yeast-GEM and rhto-GEM models used here indeed use the same style of identifiers.

### 4.6 Draft model with getModelFromHomology

Up to ten input arguments can be specified for the `getModelFromHomology` function. The first three correspond to: a cell structure containing the template model(s), the bidirectional BLAST structure, and a character string containing the organism identifier for the target organism. Note that this target organism identifier *must* match the target organism identifier provided in the input arguments of the `getBlast` function. The fourth input argument, left intentionally blank for the default value, corresponds to the prefered order of template models from which to add reactions. Here we prefer the more mature *S. cerevisiae* model. The following two input arguments correspond to options that tweak the criteria for adding reactions from the template model to the target model, which are discussed in detail in the documentation of this function. The seventh, eight and ninth argument indicate cut offs for BLAST results (maximum E-value, minimum length of alignment, and minimum identity), to inform which protein matches are likely homologs. We set a minimum alignment length of 150 bp, which is less strict than the default of 200, and will therefore result in the identification of more homologs. The default values are set as a reasonable balance between sensitivity and specificity.

### 4.7 Use expression data to adjust biomass compositions

When available, gene expression data (i.e. transcriptomics and proteomics) can be used to determine the ratio of nucleotides and amino acids in RNA and DNA, in contrast to defining these ratios based on the primary sequence only as is detailed in the protocol above. However, integration of the expression data will only have minor effect on the determine ratios, such that the simpler approach suffices.

### 4.8 Manual identification of gaps

A set of RAVEN functions can be used for identification of gaps in the draft model, which can be informative if automated approaches fail to resolve all gaps. `canProduce` and `canConsume` are functions that generate a list of metabolites that can have net synthesis or consumption, absence of biomass macromolecules from this list indicates a gap in the network. `checkProduction` calculates the smallest set of metabolites that must have net synthesis to support synthesis of all other metabolites, suggesting how gaps can most efficiently be filled. `getAllSubGraphs` can identify if there are subnetworks that are disconnected from the main metabolic network, while `haveFlux` provides a list of reactions that can or cannot carry a flux, both indicative of gaps in the network.

## Notes

https://github.com/SysBioChalmers/hanpo-GEM

https://github.com/SysBioChalmers/RAVEN

### Competing Interest Statement

The authors have declared no competing interest.

